# The *Streptomyces* volatile 3-octanone alters auxin/cytokinin and growth in *Arabidopsis thaliana* via the gene family *KISS ME DEADLY*

**DOI:** 10.1101/2020.02.15.949685

**Authors:** Bradley R. Dotson, Vasiliki Verschut, Klas Flärdh, Paul G. Becher, Allan G. Rasmusson

**Affiliations:** Lund University Department of Biology, Molecular Plant Biology Group, Lund Sweden; Swedish University of Agricultural Sciences, Department of Plant Protection Biology, Alnarp Sweden; Lund University, Department of Biology, Microbiology Group, Lund Sweden

**Keywords:** Bacterial Volatiles, 3-octanone, Plant Growth, *KISS ME DEADLY*, Auxin/Cytokinin homeostasis, *Arabidopsis thaliana*, *Streptomyces coelicolor*, *Streptomyces venezuelae*

## Abstract

Plants enhance their growth in the presence of particular soil bacteria due to volatile compounds affecting the homeostasis of plant growth hormones. However, the mechanisms of volatile compound signaling and plant perception has been unclear. This study identifies the bioactive volatile 3-octanone as a plant growth stimulating volatile, constitutively emitted by the soil bacterium *Streptomyces coelicolor* grown on a rich medium. When 3-octanone is applied to developing *Arabidopsis thaliana* seedlings, a family-wide induction of the Kelch-repeat F-box genes known as *KISS ME DEADLY* (*KMD*) subsequently alters auxin/cytokinin homeostasis to promote the growth of lateral roots and inhibit the primary root. Loss of function of the *KMD* family or other alterations of auxin/cytokinin homeostasis suppresses the volatile-induced growth response. This reveals a function of *KMD*s in the pathway of microbial volatile perception and plant growth responses.

**Significance Statement:** Volatiles from soil microbes are profound stimulators of plant growth. This work identifies for the first time a plant hormone signaling regulator, the gene family *KISS ME DEADLY* (*KMD*), to be an early essential step in plant growth promotion by a soil bacterial volatile, 3-octanone. The *KMD*-regulated gene network alters the tissue sensitivity balance for the growth hormones auxin and cytokinin, modifying root growth rate and architecture. Previously, the Kelch repeat F-box gene family of *KMD*s have been shown to be important down-regulators of both positive cytokinin signaling and phenylpropanoid biosynthesis, but upstream cues were unknown. This report places the *KMD* family regulation of plant growth and defense into its biotic context.

## Introduction

Most significant decisions that plants make are related to growth and resource allocation, such as germination (1) or shade-avoidance (2). An example of such decisions is the allocation of resources between defense and growth during plant development. Production of defensive compounds is required due to the sessile nature of plants; however, the metabolic cost depletes energy resources that could otherwise support growth and fecundity (3). Conversely, a decision to increase growth is necessarily tied to benefits that the plant can perceive in the local environment. To illustrate, during split root growth experiments with pea plants (*Pisum sativum*), the plant choice of increased root growth was found to be based on an assessment of available nutrients in the local and extended environment. Evaluation of the pea’s integrated root growth choice strategy ranks as one of the best systems for risk assessment ever tested, including man (4).

Tissue growth occurs through cell division and expansion, both of which are governed by the plant growth hormones auxin and cytokinin as well as the crosstalk between their signaling pathways (5). Direct microbial intervention in the plant hormonal homeostasis have been previously observed (6). However, recently indirect interaction via non-hormonal volatile emissions from soil microbes have been found to influence plant growth regulation (7). For example, the volatiles 2,3-butanediol and acetoin from the soil bacteria *Bacillus* spp. enhanced growth of both *Arabidopsis thaliana* and tomato (8, 9). The effect was linked to auxin signaling as the volatile-induced transcriptional responses in *A. thaliana* included an abundance of upregulated auxin signaling and cell wall remodeling genes, and the growth enhancement could be blocked by auxin transport inhibitors (8). However, developmental timing of both the plant and the microbe can influence plant responses. For instance, plant growth was enhanced in response to the volatiles from established cultures of the soil filamentous fungus *Trichoderma atroviride*, though volatiles from younger cultures of the same fungus caused growth inhibition in young plants and enhanced defense responses in older plants (10, 11).

Currently, knowledge on plant volatile perception and regulation regarding growth and/or defense is lacking with regards to the plant signaling components involved. Especially, volatile profiles from cultured soil microbes are highly diverse, and their plant growth effects depend on both the microbial strain and the culture conditions (12, 13). To this effect, we here document the previously uncharacterized role of a small family of Kelch repeat F-box proteins known as *KISS-ME-DEADLY* (*KMD*) (14) in the plant perception of volatiles from the common soil bacterium *Streptomyces coelicolor*. We identify the bacterial volatile 3-octanone as the bioactive component and dissect the affected plant gene products that influence auxin/cytokinin homeostasis and promote tissue growth.

## Results

### *Streptomyces* volatiles enhance tissue expansion and modify root architecture in *A. thaliana* seedlings

Seven-day-old *A. thaliana* Col-0 seedlings were exposed for six days to volatiles from cultures of the *S. coelicolor* M145 strain (Fig. 1a-b). Treated seedlings showed significant fresh weight increases in both roots (65%) and shoots (63%) at the highest exposure, and the growth effect was linearly correlated to *S. coelicolor* dosage (Fig. 1c). Compared to control, volatile-exposed seedlings possessed more expanded leaves, elongated petioles, shorter primary roots, and elongated lateral roots (Fig. 1a-b). Quantitative analysis of root growth rate over two days of exposure to *S. coelicolor* volatiles confirmed that the primary and lateral roots responded oppositely to the volatile communication (Fig. 1d-f, S1). The lateral root growth (LRG) increased to a rate roughly two-fold faster than control (Fig. 1d), while primary root growth (PRG) was strongly repressed compared to the untreated control (Fig. 1e). The ratio of primary to lateral root growth log2 (PRG/LRG) revealed an apparent shift in resource allocation from primary root growth to lateral root tissue investment in response to the treatment (Fig. 1f). However, the rate of lateral root emergence (LRE) was unaffected by the treatment with *Streptomyces* volatiles (Fig. 1g). These results indicate that the increased root mass observed can be attributed to a lateral root tissue expansion rather than to an increased number of organs formed.

**Figure 1.**
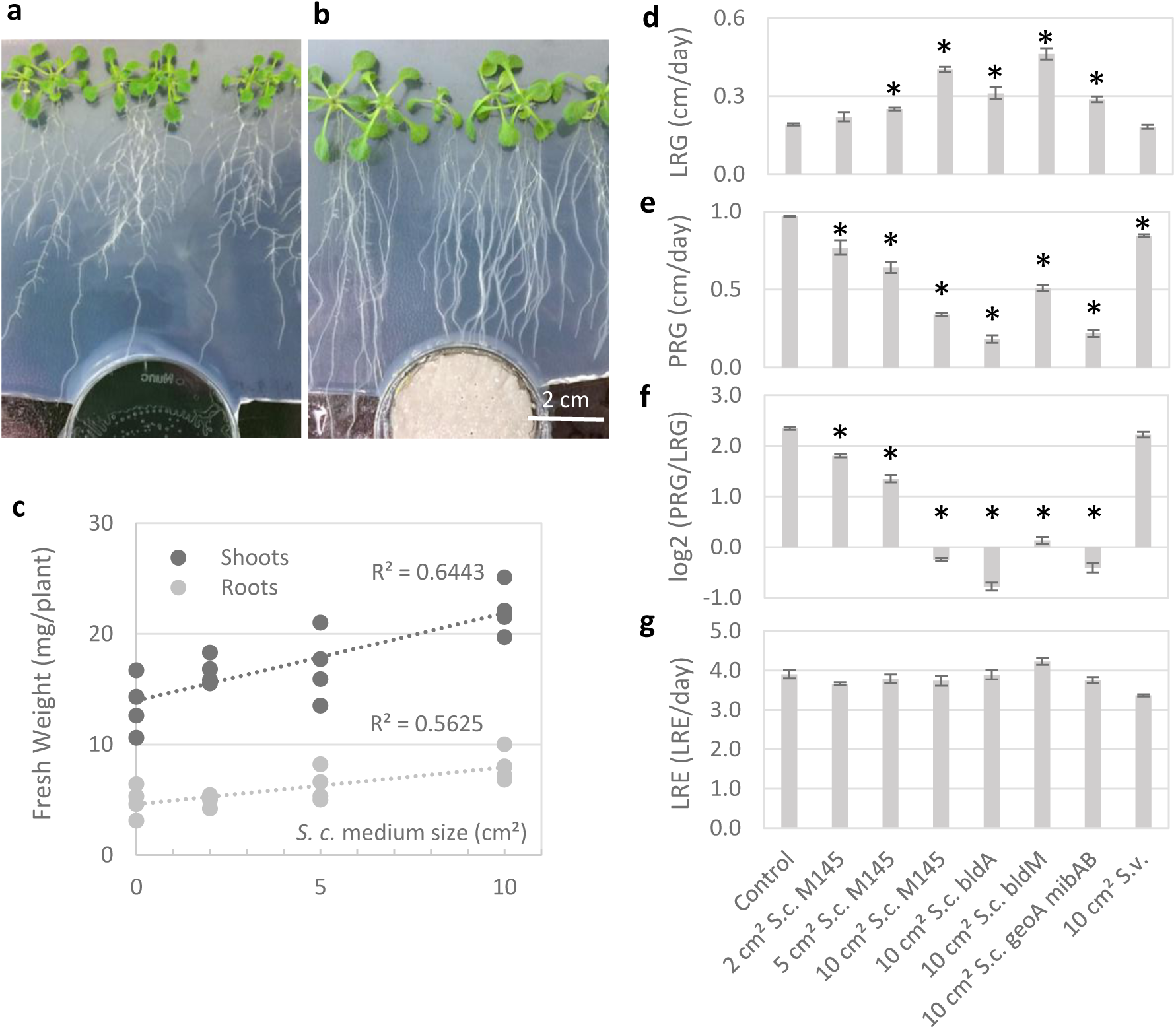
Growth of *A. thaliana* Col-0 in response to *S. coelicolor* volatiles. **a-b**, Representative images are shown for 7-day-old *A. thaliana* Col-0 seedlings that have been treated for another six days with control plates lacking bacteria (**a**) or with gas-phase contact to plates carrying *S. coelicolor* M145 grown on 10 cm^2^ of SFM (**b**). **c**, Fresh weight responses of shoots and roots of 7-day-old Col-0 that have been treated ± volatiles for an additional six days with linear regression analysis *R*^*2*^ values. Dosage of volatiles is expressed as the surface area of bacterial culture. **d-g**, Analysis of root growth responses of 7-day-old Col-0 seedlings ± exposure for two days. **d**, Lateral Root Growth (LRG) is the average growth rate of the lateral root (cm per day). **e**, Primary Root Growth (PRG) is the average growth rate of the primary root (cm per day). **f**, Log2 (PRG/LRG) is the log2 ratio of PRG to LRG. **g**, Lateral Root Emergence Rate (LRE) is the number of lateral roots emerged from the primary root per day. Bars represent standard error and asterisks indicate significant differences to the control treatment according to Student’s *t*-test with false discovery rate (FDR) correction for *q* = 0.05.

### *A. thaliana* growth is modified by volatiles from *S. coelicolor* independent of bacterial developmental stage

Volatile profiles change during development, and therefore it was tested whether the volatiles affecting *A. thaliana* seedlings are under developmental regulation in *S. coelicolor* and whether the two major volatiles geosmin and 2-methyl isoborneol, influence the plants. Cultures of *S. coelicolor* mutants of *bldA* (stalled in developmental progression to aerial mycelium formation and sporulation due to lacking an early developmental regulator), *bldM* (stalled in developmental progression to aerial mycelium formation and sporulation due to lacking a later developmental regulator), and *geoA mibAB* (lacking the major volatiles geosmin and 2-methylisoborneol), were grown on the sporulation agar medium SFM (soya flour and mannitol) and used to treat Col-0 seedlings via gas phase. All *S. coelicolor* genotypes induced similar changes in growth rate and root architecture (Fig. 1d-g, S1, S2a-b, d-f). Likewise, the changes in root architecture occurred without regard to the presence of sucrose in the plant medium (S2g-h), indicating that plant internal metabolic resources were not active determinants in our assays.

### Screening of volatiles from different *Streptomyces* strains

To identify the compound or compounds stimulating *A. thaliana* growth, we analyzed the correlation between volatile profiles from different *Streptomyces* strains and their effects on plant seedlings. Analysed cultures included *S. coelicolor* strains M145, *bldA* and *bldM*, as well as *S. venezuelae*, which when cultivated on its regular sporulation medium (MYM; maltose, yeast extract, and malt extract), resulted in little or no volatile effect on the growth of *A. thaliana* seedlings (Fig. 1d-g, S1, S2c). Volatiles were collected from the headspaces of *Streptomyces* cultures for a period of 24 h and analyzed using GC-MS (Table 1). The profiles of volatile compounds were evaluated to identify compounds exclusively emitted by the *Streptomyces* cultures that induce the root growth responses (Table 1). Overall, the compounds identified were organic, small and hydrophobic. The volatile compounds 3-octanone and two isomers of chalcogran were exclusively found in the headspace of the plant growth-inducing *S. coelicolor* cultures. The presence of 3-octanone and chalcogran in growth-stimulating headspace indicated that at least one of these compounds induced growth stimulation (Table 1). The compounds acetoin and 2, 3 butanediol were specifically targeted as they have previously been found to have growth effects on *A. thaliana* when released from bacteria (15). However, these compounds were undetectable in all *Streptomyces* headspaces. Most of the other compounds identified have previously been observed among volatiles from various soil microbial species (12).

**Table 1.** Volatila compounds identified in *Streptomyces* headspace. Volatile headspace from four different *Streptomyces spp*. genotypes grown on sporulation media was analyzed by GC-MS using a DB-Wax column. Volatile compound name, percent of total volatile headspace ± standard error, Kovats’ index, CAS number and method of identification is listed from each of the *Streptomyces* headspace analyses. The compounds that were identified specifically in the headspaces of growth-inducing cultures are highlighted. Identification (ID) methods are by Mass Spectrometry (MS), Kovats’ Index (KI) and coinjection of authentic reference compounds (RC). Unidentified compounds are listed by letters as well as mixed volatile profile peaks of two or more compounds are denoted by §. Data for compound quantities are average percentages of total volatile content ± standard error or not detectable (N.D.).

| Volatile compound | *Streptomyces coelicolor* M145 | *Streptomyces coelicolor* bldA | *Streptomyces coelicolor* bldM | *Streptomyces venezuelae* | Kovats' Index experimental | Kovats' Index Published | CAS number | ID Method |
| --- | --- | --- | --- | --- | --- | --- | --- | --- |
| A-unknown | 0.03 ± 0.01 | N.D. | N.D. | N.D. | 1134 | - | - | MS |
| B-unknown | N.D. | N.D. | N.D. | 6.8 ± 0.3 | 1147 | - | - | MS |
| 3-Octanone | 0.15 ± 0.02 | 2.9 ± 0.8 | 0.8 ± 0.1 | N.D. | 1260 | 1266 | 106-68-3 | MS, KI, RC |
| Styrene | 0.26 ± 0.04 | 2.0 ± 0.4 | 1.6 ± 0.2 | 0.3 ± 0.1 | 1263 | 1259 | 100-42-5 | MS, KI |
| Chalcogran (i) | 0.3 ± 0.1 | 0.7 ± 0.1 | 1.0 ± 0.3 | N.D. | 1357 | 1343 | 35401-84-2 | MS, KI, RC |
| Chalcogran (ii) | 0.4 ± 0.2 | 1.1 ± 0.1 | 1.3 ± 0.4 | N.D. | 1361 | 1348 | 35401-84-2 | MS, KI, RC |
| D§-unknown | 0.36 ± 0.04 | N.D. | 1.3 ± 0.3 | N.D. | 1436 | - | - | - |
| 2-Methyl-isoborneol | 6.4 ± 0.8 | 3.2 ± 0.3 | N.D. | 60.7 ± 4.6 | 1595 | 1602 | 2371-42-8 | MS, KI, RC |
| E unknown | 0.33 ± 0.04 | N.D. | 2.7 ± 0.4 | N.D. | 1610 | - | - | MS |
| F unknown | N.D. | 0.7 ± 0.4 | N.D. | 1.7 ± 1.6 | 1714 | - | - | MS |
| Germacrene D | 9.9 ± 1.9 | 0.9 ± 0.4 | N.D. | 1.2 ± 0.1 | 1724 | 1724 | 483-76-1 | MS, KI |
| G unknown | N.D. | 42.8 ± 3.1 | 30.8 ± 2.4 | N.D. | 1732 | - | - | MS |
| H unknown | N.D. | N.D. | N.D. | 1.0 ± 0.7 | 1747 | - | - | MS |
| Geosmin | 49.1 ± 3.2 | 6.9 ± 0.2 | N.D. | 6.2 ± 2.0 | 1845 | 1858 | 19700-21-1 | MS, KI, RC |
| I-unknown | 1.7 ± 0.5 | N.D. | N.D. | 4.3 ± 0.4 | 2150 | - | - | MS |

### *A. thaliana* hormone mutants have altered responses to *S. coelicolor* volatiles

Mutants for central signaling genes in several hormone pathways of *A. thaliana* were screened to identify signaling processes involved in the plant response to *S. coelicolor* volatiles. For each plant genotype, we determined the log2 root growth rate ratio for presence and absence of *S. coelicolor* M145 volatiles, separately for lateral root growth (Δ log2 LRG) and primary root growth (Δ log2 PRG). This allowed normalization of the responses in the phenotypically diverse sets of mutants (Fig. 2). Exposure of Col-0 to the volatiles caused a 2-fold increase in lateral root growth rate, and a 2.8-fold decrease in primary root growth rate during 2 days of treatment. In contrast, auxin and cytokinin response mutants, including the cytokinin-resistant mutant *cre1-12*, the auxin transcription factor *arf1-2*, the auxin influx transporter *aux1-7* and the F-box receptor for auxin *tir1-1*, were overall unaffected by the volatiles after 2 days of exposure. However, observation of morphology after 6 days (S3), indicated that the growth response in *tir1-1* may be merely delayed. Unlike the loss of sensitivity in the cytokinin and auxin mutants, the ethylene signaling mutant *ein2-1* lacked the lateral root growth increase of Col-0, but the primary root inhibition was retained. Additionally, the strigolactone-insensitive mutant *max2-1* showed less pronounced reductions in both primary and lateral root growth effects. Hormone mutants for other pathways, *abi1-1, gai-1, coi1-37*, and *bri1-4* were also tested, but had mild to no discernible reductions in the volatile-induced growth effects (S3). Further verification of auxin involvement was carried out with the auxin response reporter DR5::GFP, which showed alterations of auxin distribution in the root tip upon treatment with *S. coelicolor* volatiles (S4). The results thus clearly indicate that the primary and lateral root growth responses to *S. coelicolor* volatiles involve the auxin and cytokinin response pathways.

**Figure 2.**
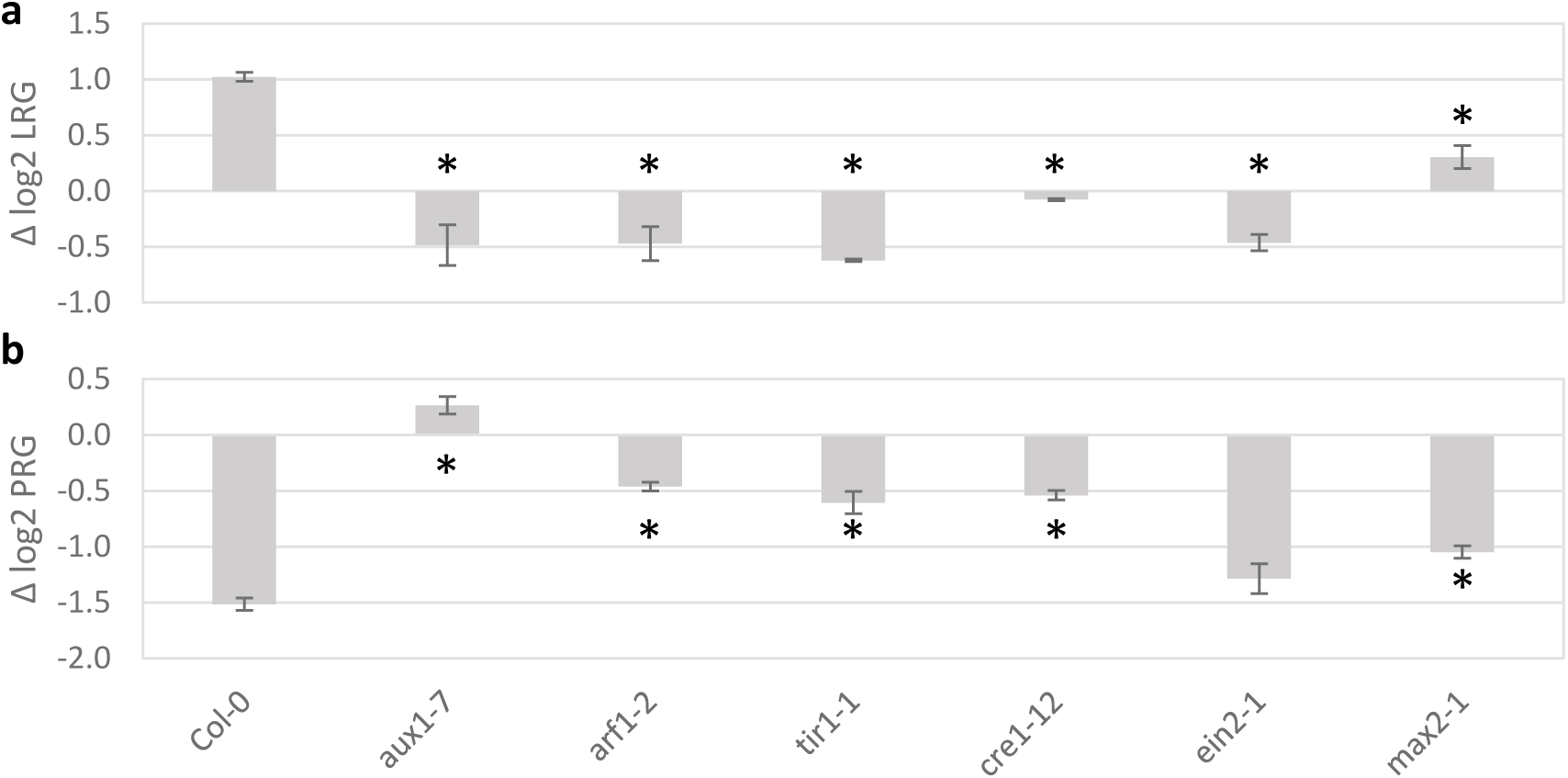
Growth of *A. thaliana* wild-type and hormone mutants in response to *S. coelicolor* volatiles. Seven-day-old seedlings of wild-type and hormone mutants were treated for two days with volatiles from *S. coelicolor* M145 grown on 10 cm^2^ of SFM sporulation medium. The figure shows log2 ratios of growth ± volatiles for primary (**a**; Δ log2 LRG) and lateral (**b**; Δ log2 PRG) roots. Bars represent standard error and asterisks indicate significant differences to Col-0 according to Student’s *t*-test with FDR correction for *q* = 0.05.

### The *A. thaliana* response to Streptomyces volatiles shows similarity to other biotic responses

RNA was isolated from control and *S. coelicolor* volatile-treated seedlings and analysed by transcriptome sequencing (RNA-Seq). A total of 18,493 transcripts were identified, whereof 26 were significantly differentially expressed with a ratio of at least 2 fold (Dataset 1). In order to determine the biological relevance of the response to *S. coelicolor* volatiles, the RNA-Seq expression profile of the 26 significantly responsive genes was compared to public *A. thaliana* gene expression data sets using the Genevestigator signature search tool (Table 2). The highest profile similarity was found to the *A. thaliana* response to volatiles from the rhizobacterium *Serratia plymuthica* HRO-C48 (16). Other similar datasets were for root colonization by the beneficial *Ascomycota* endophyte *Colletotrichum tofieldiae* and the closely related pathogen *Colletotrichum incanum* during phosphate starvation (17). The RNA-Seq profile observed here is thus consistent with previous biotic treatments of *A. thaliana*.

**Table 2.**
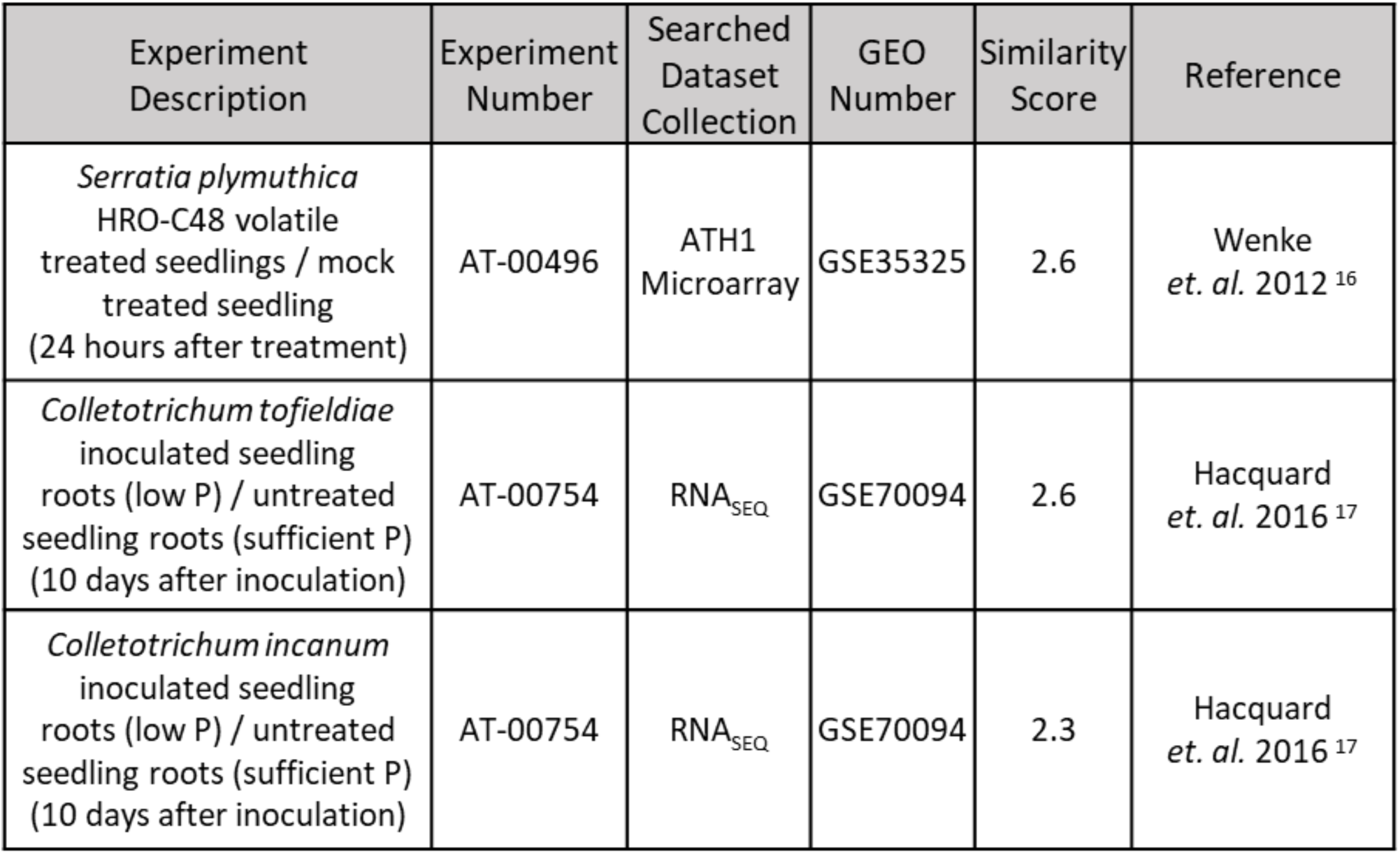
Similarity analysis of volatile-induced *A. thaliana* RNA profiles. Similarity analysis of volatile-induced *A. thaliana* RNA profiles. Seven-day-old *A. thaliana* Col-0 seedlings were exposed to a 10 cm^2^ Streptomyces culture for 2 h and analysed by RNA-Seq. RNA-Seq data profiles for genes that were found to be significantly differentially expressed from control (q < 0.05) and having a volatile response of at least 2-fold were analyzed by Genevestigator Signature Search against responses in publically available microarray and RNA-Seq *A. thaliana* gene expression datasets using Pearson correlation.

### Gene ontology analysis of RNA-Seq data identifies initial plant reprogramming components

Investigations of gene ontology (GO) was carried out by non-parametric analyses of fold changes for all 18,493 transcripts using the programs Gorilla and Genevestigator (Table 3a). The combination of analyses identified the GO bins of “Negative Regulation of Cytokinin-Activated Signaling Pathways” and “Biological Process” with the lowest FDR *q*-value, followed by “Response to Stimulus”, “Cell Wall Organization” and “Regulation of Phenylpropanoid Metabolic Process”. Two GO bins found, “Regulation of Phenylpropanoid Metabolic Process” (Gorilla) and “Negative Regulation of Cytokinin-Activated Signaling Pathways” (Genevestigator) are both highly enriched in a small family of kelch repeat F-box genes *KISS ME DEADLY* (*KMD1-4*) (Table 3b, Dataset 2). Moreover, the “Cell Wall Organization” GO bin mainly contain pectin degradation and extensin-like proteins typically involved in cell wall loosening associated with growth changes (18).

**Table 3.** Gene ontology analysis of *S. coelicolor* volatiles-induced RNAs. **a**, Gene Ontology analysis was performed on the total list of 18,493 genes, sorted by fold changes. Resulting GO terms, bin sizes, and descriptions of FDR *q*-values are as provided by Gorilla and Genevestigator, and descriptions are as annotated by AmiGO. All bins shown had primary *p*-values < 0.001. **b**, Description of the *KISS ME DEADLY* family of genes identified by GO analyses as a significant group of genes in bins of GO:0080037 and GO:2000762. Fragments per kilobase of exon model per million reads mapped (FPKM) and the complete data set can be found in Dataset 1.

| Program | GO Term | GO Bin Hits /<br>GO Bin Size | Description of Bin | FDR<br>*q*-value |
| --- | --- | --- | --- | --- |
| Genevestigator | 0080037 | 3/3 | Negative Regulation of Cytokinin-Activated Signaling Pathways | 0.008 |
| Genevestigator | 0008150 | 116/22565 | Biological Process | 0.008 |
| Genevestigator | 0050896 | 52/6661 | Response to Stimulus | 0.014 |
| Gorilla | 0071555 | 20/109 | Cell Wall Organization | 0.071 |
| Gorilla | 2000762 | 4/5 | Regulation of Phenylpropanoid<br>Metabolic Process | 0.090 |

**b**
| Gene<br>Number | Gene<br>Description | Average<br>FPKM;<br>Control | Average<br>FPKM;<br>*S.c.*<br>Volatiles | Fold Change;<br>*S.c.* Volatiles<br>/ Control | GO Bin |
| --- | --- | --- | --- | --- | --- |
| At1g80440 | Galactose oxidase/kelch<br>repeat superfamily protein;<br>*KISS ME DEADLY 1* | 95.8 | 182.5 | 1.9 | GO:80037;<br>GO:0008150;<br>GO:0050896;<br>GO:2000762 |
| At1g15670 | Galactose oxidase/kelch<br>repeat superfamily protein;<br>*KISS ME DEADLY 2* | 104.1 | 171.2 | 1.7 | GO:80037;<br>GO:0008150;<br>GO:0050896;<br>GO:2000762 |
| At2g44130 | Galactose oxidase/kelch<br>repeat superfamily protein;<br>*KISS ME DEADLY 3* | 67.7 | 113.2 | 1.7 | GO:0008150;<br>GO:0050896;<br>GO:2000762 |
| At3g59940 | Galactose oxidase/kelch<br>repeat superfamily protein;<br>*KISS ME DEADLY 4* | 221.4 | 302.6 | 1.4 | GO:80037;<br>GO:0008150;<br>GO:0050896;<br>GO:2000762 |

### Transgenic plants deficient in KMD and associated cytokinin response proteins are unresponsive to *S. coelicolor* volatiles

The KMD protein family consists of four regulatory F-box proteins that target cytokinin-responsive Type B-ARR transcription factors (ARR1, 10 and 12) for ubiquitination-mediated degradation (14, 19). In addition, the KMD proteins target a set of proteins that function in the initiation of the phenylpropanoid biosynthesis pathway (20, 21). We found that the *A. thaliana* triple mutants *kmd1,2,4* and *kmd1,2,α4* had suppressed lateral root growth increases, and primary root growth inhibitions in response to *S. coelicolor* volatiles (Fig. 3, S4). Additionally, the cytokinin-insensitive triple mutant of the Type-B-ARR genes *arr1,10,12* displayed enhanced sensitivity in the primary root and lacked the volatile-induced increase in lateral root growth. Further, repressor factors of the cytokinin response known as type-A ARR proteins, were tested using quadruple mutants *arr3,4,5,6* and *arr5,6,8,9*. Both type-A ARR lines displayed volatile-resistant phenotypes, the *arr3,4,5,6* being especially resistant to primary root inhibition, and *arr5,6,8,9* to lateral root stimulation, by *S. coelicolor* volatiles.

**Figure 3.**
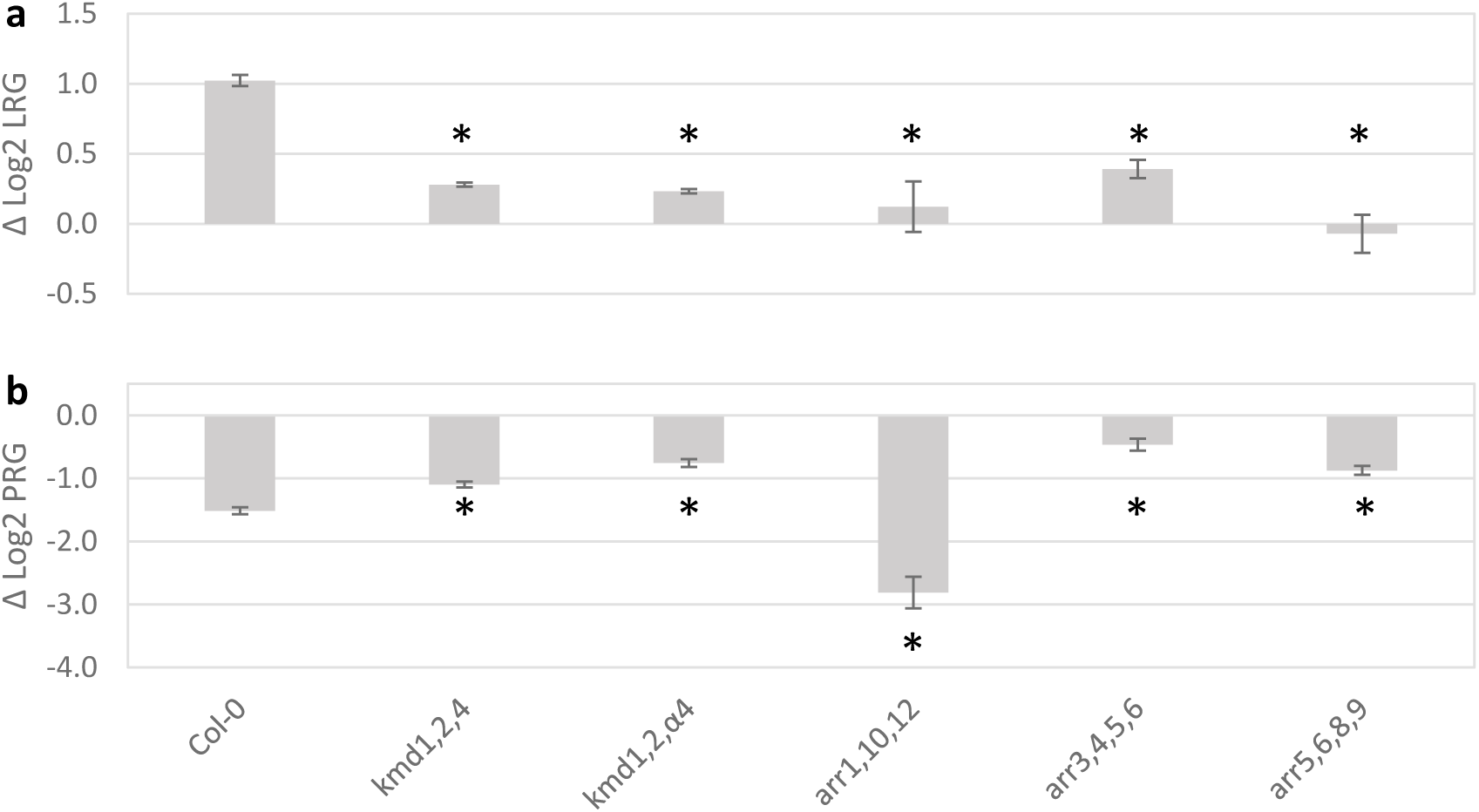
Volatile responses of mutants for *KMD* and associated hormone response genes. The figure shows log2 ratios of growth ± volatiles for lateral roots (**a**; Δ log2 LRG) and primary roots (**b**; Δ log2 PRG) of Col-0, *KISS ME DEADLY* and related hormone mutants. Seven-day-old seedlings were exposed for two days to volatiles from *S. coelicolor* on 10 cm^2^ of SFM. Bars represent standard error and asterisks indicate significant differences to Col-0 according to Student’s *t*-test with FDR correction for *q* = 0.05.

### The root growth response elicited by the bioactive volatile 3-octanone depends on *KMD* and auxin/cytokinin homeostasis

In order to directly test pure volatiles and KMD for involvement in the response to *S. coelicolor*, 7-day-old seedlings were transferred to agar plates containing pure compound or solvent control and lateral and primary root growth was quantified over 2 days (Fig. 4). Initial studies of Col-0 seedlings with 50 µM chalcogran showed limited effects and were not elaborated on further (Fig 4c). In contrast, Col-0 treated with 3-octanone displayed dose-dependent inhibition of the primary root and stimulation of the lateral root (Fig. 4), as observed for total *S. coelicolor* volatiles (Fig. 1). Thus 3-octanone induces a 3-fold relative growth rate shift, from primary to lateral roots (Fig. 4). We further tested mutants lacking *KMD* and other signaling components of cytokinin and auxin responses. All were resistant to the 3-octanone effect on lateral and primary root growth (Fig. 4d-e). The lack of effects, being in stark opposition to the 3-octanone-induced root growth changes in Col-0, indicates that the integral network of *KMD* as well as cytokinin and auxin homeostasis, must be intact for *S. coelicolor* volatiles or 3-octanone-induced responses to occur.

**Figure 4.**
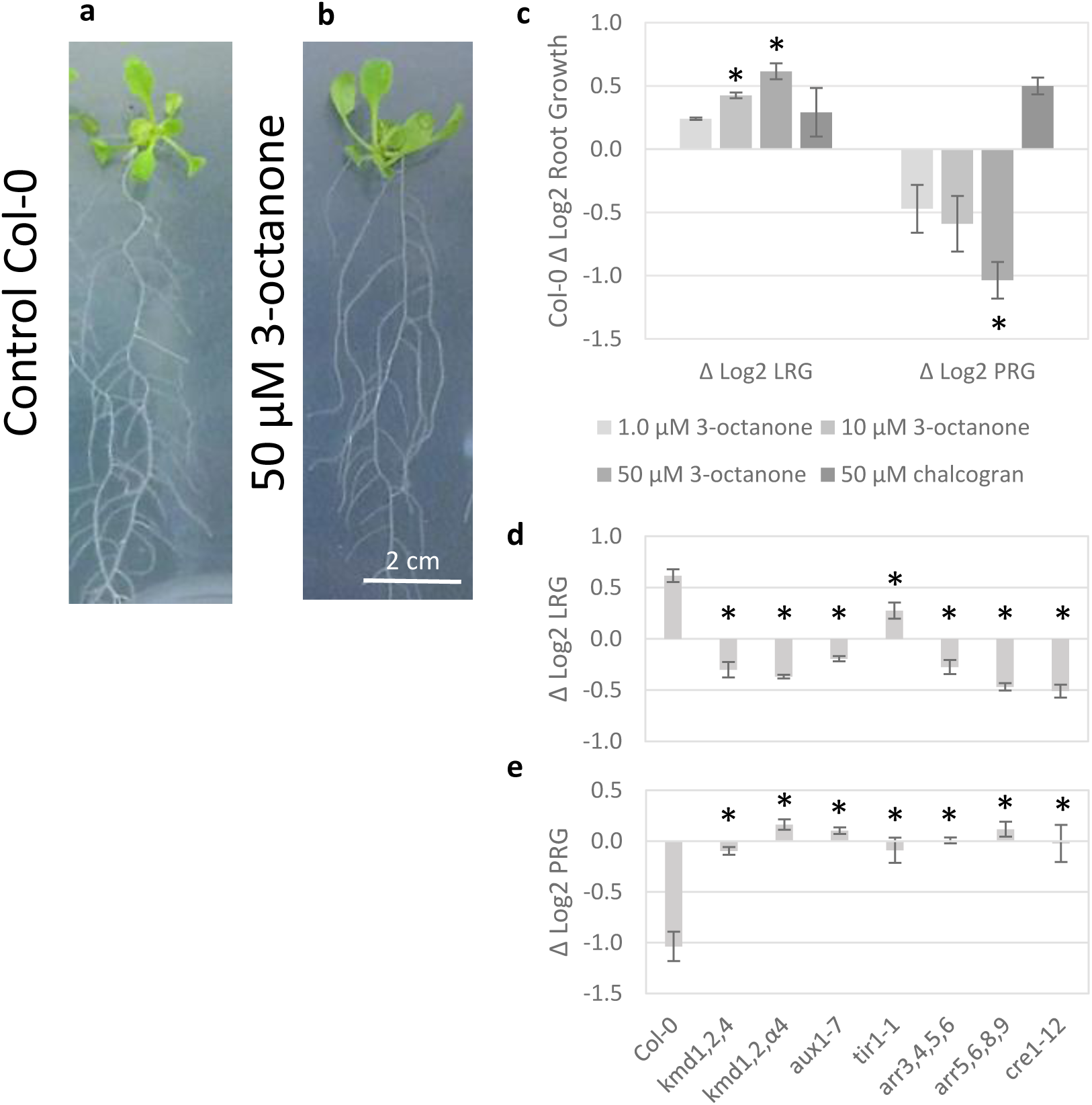
Root growth responses to 3-octanone. Seven-day-old seedlings were transferred to media containing different concentrations of 3-octanone or chalcogran. Enhanced growth of roots and shoots can be seen in representative seedlings treated on media containing 3-octanone or solvent control for six days (**a** and **b**). Root growth was quantified during two days of treatment: **c**, shows log2 growth ratios ± volatile compound (volatile-treated / control) for lateral (Δ log2 LRG) and primary (Δ log2 PRG) roots. **d-e**, show comparisons of Col-0 and signaling mutants for log2 growth rate ratio ± 50 µM 3-octanone for lateral (**d**) and primary (**e**) roots. Bars represent standard error and asterisks indicate significant differences between growth ± volatiles (**c**) and to Col-0 (**d-e**) according to Student’s *t*-test with FDR correction for *q* = 0.05.

## Discussion

*A. thaliana* is adapted to an extensive range of nutrient availability and has consequently evolved decision mechanisms to adapt to variable local soil environments (22, 23). Root growth investments in soil niches with high microbial activity have previously been shown to improve plant overall growth by enhancing plant nutrient status, pathogen resistance and by mitigating stress responses (24, 25). It is therefore expected that higher levels of microbial activity would be cues for plant decisions to locally enhance root tissue investments (26). We identified 3-octanone as a bio-active volatile signal that is produced by *S. coelicolor* grown on SFM sporulation media. This happens regardless of the bacterial development, because *S. coelicolor* mutants stalled at different developmental states still emit the volatile 3-octanone and influence the *A. thaliana* root growth (Fig. 1d-f, Table 1).

The 3-octanone volatile has previously been shown to be one of many compounds released from *in vitro Streptomyces* cultures(27), but has not been shown to be produced by vascular plants. Along with 1-octen-3-ol, 3-octanone belongs to a class of eight-carbon (C8) compounds that possess bioactivity in organisms like bacteria, fungi, and plants. Notably, 3-octanone is a conidiation-inducing signal in the fungal genus *Trichoderma* (28), which includes several plant-symbiotic species (29). *Trichoderma*-produced 3-octanone have been shown to promote growth of plants such as *A. thaliana* and willow (30, 31). In contrast, 1-octen-3-ol induces defense responses in *A. thaliana*, suppresses growth and development of fungi and reduces growth, chlorophyll accumulation and seed germination in *A. thaliana* (32–34). The bioactive C8 compounds share structural similarities with the defense regulators C9 oxylipins, both being derived from linoleic acid (35). Unlike the C9 oxylipins, there are currently no known components mediating the perception or response to C8 compounds in higher plants. Therefore our data present the first evidence linking C8 volatiles with *KMD* expression and root growth modulation.

Volatiles from *S. coelicolor* cultured on the *S. venezuelae* sporulation medium for (MYM) had little effect on *A. thaliana* root growth, whereas *S. venezuelae* enhanced lateral root growth only when cultured on *S. coelicolor* sporulation medium (SFM; S2i-j). This indicates that the *Streptomyces* medium is a critical factor for volatile plant growth promotion, rather than the *Streptomyces* species. Likewise, *in vitro* studies of several other rhizospheric bacteria have found that plant growth promotion by volatiles depends on the culture media used for the microbe (12). The two compounds correlating with plant growth effects in our assays, 3-octanone, and chalcogran have been identified as signature volatiles from fungal cultures on media high in linoleic acid (36, 37). This fatty acid is highly abundant in soya flour, which is a main component of the SFM medium (∼1.4g linoleic acid per L) (38, 39). Unsaturated fatty acids, including linoleic, tend to be degraded rapidly in soil by abiotic conditions and by microbes (40). To plants, soils that are enriched in linoleic acid-associated microbial volatiles may indicate a richness in organic compounds and availability of new nutritional resources for plant roots to harvest.

To investigate factors regulating the *A. thaliana* response to volatiles from *S. coelicolor*, exposed seedlings were analysed by RNA-Seq and displayed a differential expression pattern that matched previously shown *A. thaliana* responses to microbials (Table 2) (16, 17). Two GO approaches specifically indicated the gene family *KISS-ME-DEADLY* (*KMD*) as contributors to the *S. coelicolor* volatile-induced gene response profile in *A. thaliana* (Table 3). The GO bin with lowest FDR *q-*value, “negative regulation of cytokinin-activated signaling pathways”, contains three out of the four genes in the *KMD* family, providing an association with suppression of type-B ARR cytokinin signaling factors (14). Additionally, the entire family of *KMD* genes are included in the bin “Regulation of phenylpropanoid metabolic process” denoting the association of *KMD* genes with suppression of the phenylpropanoid biosynthesis pathway (Table 3) (20).

Concurring with transcript analysis, investigations into broad hormone regulation by *S. coelicolor* volatiles response was analyzed by genetic perturbations in the plant hormone pathways. Mutant lines impaired in auxin response (*aux1-7, arf1-2*, and *tir1-1*) and cytokinin (*cre1-12*) signaling had the smallest initial root growth response to *S. coelicolor* volatiles (Fig. 2). Consistent with the GO analysis, mutations to KMD family function suppressed both primary and lateral root responses to *S. coelicolor* volatiles as well as to the pure 3-octanone (Fig. 3). The mutant lacking the targets of KMD function, Type B ARRs (ARR1,10,12), also lacked lateral root growth enhancement but retained a strong primary root inhibition, possibly due to KMD suppressing remaining type B ARRs, such as ARR20 (14). Mutations in the cytokinin response-suppressive factors Type A ARRs (*arr3,4,5,6* and *arr5,6,8,9*) also suppressed the *S. coelicolor* volatile growth response further emphasizing that repression of cytokinin signaling is involved in mediating the *S. coelicolor* volatile-induced plant growth decision. Previous research has characterized the downstream signaling effects of KMD, which causes a shift in the cytokinin to auxin homeostasis toward more auxin-like responses (e.g., increased cell expansion and enhanced lateral root growth) and promotes synthesis of defense-associated phenylpropanoids (14, 19, 41). In this study, we identify an upstream cue for the *KMD* family, which therefore can be placed as a central auxin/cytokinin crosstalk regulator of root growth plasticity in response to a biotic interaction. We suggest a model for the consecutive steps from KMD activation, via changes in auxin to cytokinin ratios that promote wall loosening and cell expansion (5, 18) (Fig. 5). The model is consistent with known cross-talk events that balance the auxin/cytokinin homeostasis. One such example involves *SHORT HYPOCOTYL2* (*SHY2*), which is induced by positive regulators of cytokinin signaling of the Type-B *ARR* family (14, 19). *SHY2* suppresses PIN-mediated auxin membrane transport, causing a local auxin accumulation in cells with high amounts of cytokinin (Fig. 5)(42). When auxin accumulates, degradation of SHY2 via the SCF^TIR1^ ubiquitination pathway and promotion of *AHP6* by ARF5 decreases the cytokinin suppression of auxin transport (5), thus mediating a developmentally tight control of growth. KMD family proteins induce degradation of Type-B ARR 1,10,20 proteins (Fig. 5) (14, 19), so inactivation of the *KMD gene*s leads to accumulation of Type-B ARRs and cytokinin hypersensitivity (14). A volatile-induced KMD-mediated loss of cytokinin signaling would decrease the suppression of auxin signaling (e.g. via the *ARR1,10-*dependent auxin-suppressing gene *SHY2*) and enhance *PIN* expression and tissue elongation (42). Supporting such a scenario for the root growth effect of *S. coelicolor* volatiles, the GO analysis also indicated induction of genes for cell wall loosening enzymes (Fig. 5, Dataset 2), which are associated with cell expansion (18). The lack of transcriptional differences between control and volatile-treated plants for positive auxin signaling, metabolism, or transport further indicates that the observed promotion of lateral roots is a result of *S. coelicolor* volatiles decreasing cytokinin-induced suppression of auxin signaling.

**Figure 5.**
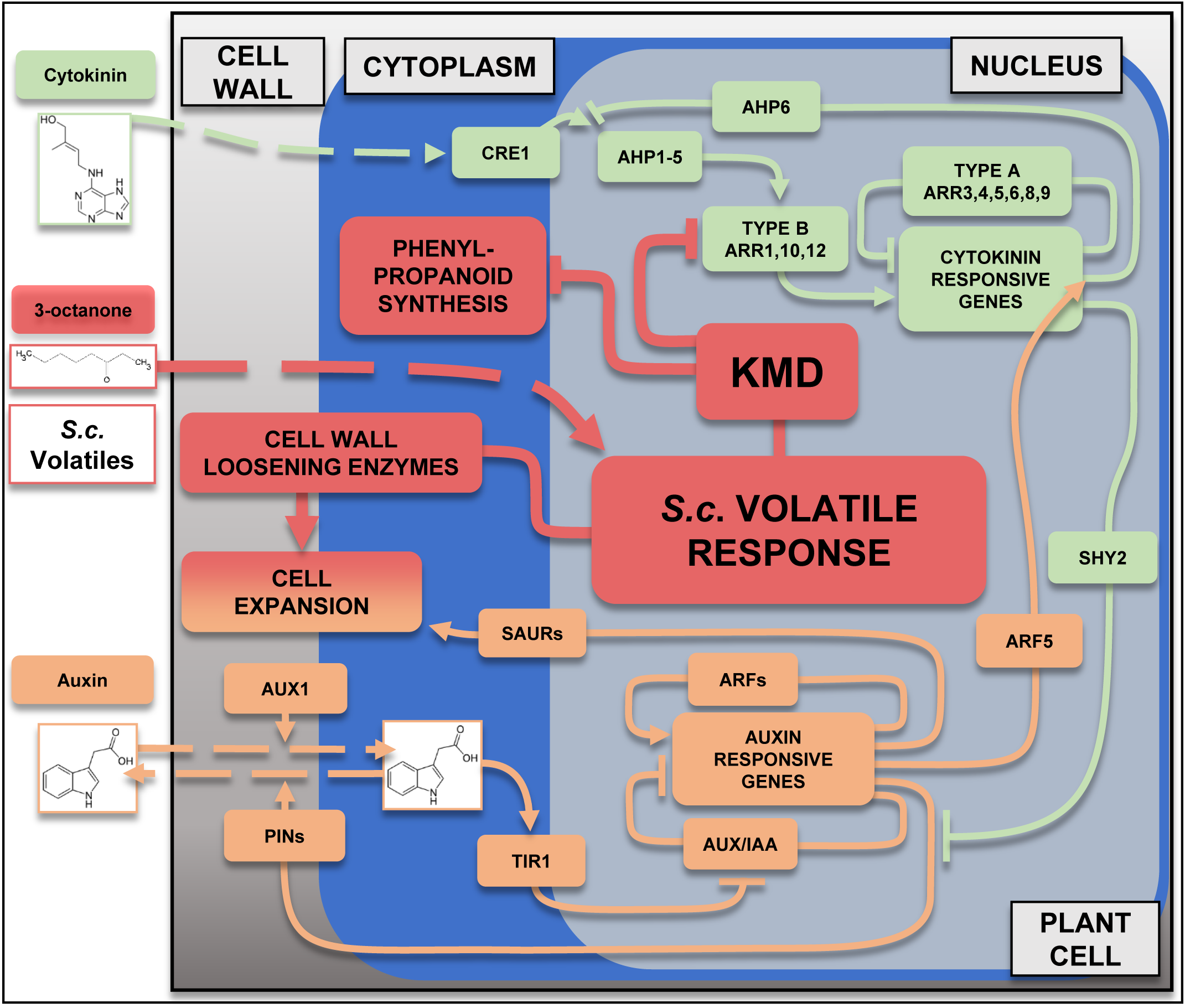
A model for *S. coelicolor* volatile induction via KMD proteins at the auxin-cytokinin intersection. Components induced by *S. coelicolor* (*S*.*c*.) volatiles are colored in red, auxin pathway components in orange and cytokinin response pathway components are in green. Arrows ending in arrows or blocks indicate functions in activation or suppression respectively. The *A. thaliana* volatile response is caused by 3-octanone perception by the plant cell. This leads to the upregulation of the KISS ME DEADLY (KMD) family of proteins that suppress cytokinin signaling through the degradation of Type-B ARR proteins (14), as well as inhibiting phenylpropanoid metabolism (20). Auxiliary to this, pectinolytic cell wall-modifying enzymes are enhanced. The reduction of type-B ARR proteins would lead to decreased expression of *SHY2*, decreasing its suppression of the auxin efflux transporters *PIN*s, leading to increased auxin transport and signaling (5). Auxin is transported into the plant cell via the influx transporter AUX1/LAX family (5). Auxin then becomes a ligand of the F-box receptor protein TIR1 that bind and inhibit the AUX/IAAs transcriptional repressors, preventing their interaction with promoter elements in auxin inducible genes (5). ARF transcription factors are instead attracted to the promoters and promote the transcription of auxin responsive genes such as the SMALL AUXIN UP RNAs (SAURs) that promote cell expansion (43). Also induced by ARFs are the *AUX/IAA* repressing transcription factors which then act as a close valve for the system (5). ARF5 has an additive response to promote the cytokinin signaling suppressor AHP6, which inhibits the cytokinin receptor AHK/CRE1 phospho-transfer to AHP1-5 proteins (5). The reduced phosphorylation of AHP1-5 leads to inactivation of Type-B ARR proteins and enhanced suppression by ambient Type-A ARRs proteins (5).

## Methods

Details about the statistical analysis data are described in the Dataset 3. Genotypes and strains used are described in S6.

### Plant growth system

Seeds of *A. thaliana* were surface-sterilized in microtubes with 70% ethanol, then 50% bleach, and finally washed three times with diH_2_O and stratified for 2 days at 4°C (44). The seeds were then placed on vertical 12 x 12 cm 0.8% agar (w/v) plates containing ½ Murashige and Skoog (MS) (45) basal medium (pH 5.50-5.65) supplemented with 88 mM sucrose, unless otherwise stated. Agar plates supplemented with volatile compounds (such as 3-octanone) were prepared similarly, except that the compounds were added from sterile filtered stocks in dimethyl sulfoxide (DMSO) to the autoclaved growth media after allowing the media to cool to <60°C. Thus, the volatile-supplemented agar plates also contained 0.01 µM DMSO, which was also used in control plates. Prepared agar plats were used immediately. The plates with seeds were incubated in a growth chamber (16 h light period, 120 µE m^-2^ s^-1^, 25°C,).

Vertical plates with growing plants were modified by removing the bottom 4 cm of the agar medium and placing a 3.5 cm petri dish inside. Variably sized disks of sporulating *Streptomyces* cultures were placed onto the 3.5 cm mini Petri dish after 7 days of plant growth. The plant-bacterial plates were then scanned with a Ricoh aficio MP C5000 scanner every 2 days over a 6 day period to measure growth differences. After six days of exposure, the plates were photographed using an iSight camera (Apple). All image processing was done by ImageJ 1.48v (National Institute of Health) (46).

### Bacterial plates

Approximately 5 µL of a spore suspension was spread over sporulation media, which was soya flour-mannitol media (SFM) for all strains of *S. coelicolor*, and maltose-yeast extract-malt extract (MYM) media supplemented with 0.2% trace minerals for *S. venezuelae* (47). *Streptomyces* cultures were allowed to grow to a dense lawn of mycelium over a period of 8 days at 30°C before being use for volatile treatments.

### Analysis of tissue growth

Primary and lateral root growth was quantified over a period of two days by ImageJ analyses of scanned photos. Lateral root growth was assessed using the largest three existing lateral roots per seedling and primary root growth was assessed per seedling. Experimental growth rates were compared internally as the overall lateral root growth/primary root growth (log2 LRG/PRG). The number of emerging lateral root per day (LRE) was calculated per primary root present. Alternatively, volatile treatments were compared to control settings for lateral root growth (Δ log2 of volatile-treated LRG to log2 control LRG) and for primary root growth (Δ log2 of volatile-treated PRG to log2 control PRG). For each measurement of growth, averages from technical replicates of 3 or more biological replicates were pooled into 3 biological replicates and used to compare between experimental and control. The significance of growth differences was tested by Student’s *t*-test and verified with an FDR *q*-value of 0.05 according to Benjamini & Hochberg (1995) (48).

### Volatile collection and analyses

Volatiles were collected from 8-day old *Streptomyces* cultures over a period of 24 h by adsorption to air filters made of Teflon tubes (4 × 40 mm), holding 50 mg Porapak Q (80/100 mesh; Alltech, Nicholasville KY, USA). Adsorbed volatiles were desorbed by eluting from the filters with 500 µl re-distilled n-hexane. Two µL of collected volatiles were resolved by gas chromatography-mass spectrometry (GC-MS; 6890 GC and 5975 MS; Agilent Technologies Inc., Santa Clara, CA, USA) using a polar DB-WAX capillary column (30 m × 0.25 mm × 0.25 μm; J&W Scientific, Folsom, CA, USA) using helium as carrier gas at a flow rate of 35 cm / second. The oven temperature was held at 30 °C for 3 min, increased with a ramp rate of 8 °C/min to 225 °C, and held for 10 min. Compounds were then subjected to mass spectrometry in a GC-MS (6890 GC and 5975 MS; Agilent Technologies Inc., Santa Clara, CA, USA). Compound identities were determined according to mass spectra libraires at SLU Alnarp and library of the National Institute of Standards and Technology (NIST; www.nist.gov), retention times (using the Kovats’ index (49) NIST database for polar columns) and by co-injection of authentic compound standards (Sigma).

### RNA extraction

Single replicate experiments with volatile and control treatment were independently repeated three times using populations of approximately 20 seedlings. Seedlings were exposed to control or *S. coelicolor* M145 volatiles conditions for 2 h and flash frozen in liquid N_2_. Approximately 100 mg of tissue was placed into a microtube and then ground with plastic pestles in a. RNA was extracted according to the protocol of the RNeasy kit (Qiagen, Venlo, Netherlands), and treated with the DNA-*free* DNA Removal kit (ThermoFisher, Waltham, MA USA). About 1 µg of each RNA sample was sent to Beijing Genomic Institute (Hong Kong, China) for RNA-Seq analysis using the Hiseq 2000 Illumina sequencing platform and single-end 50 base pair sequencing.

### Analysis of RNA-Seq data

Raw reads were mapped to the genome of *A. thaliana* (TAIR 10) (50) as described (51). Reads were summarized per gene (including splice variants) and divided by mRNA length to calculate reads per kb million (RPKM), which were log2-transformed. Transcripts identified as non-coding or organellar, as well as lowly expressed genes (less than 15 mapped reads per gene), were removed and not being part of the analyses. Transcripts were then mean expression-normalized according to Willems *et al*. 2008 (52) for calculation of log2 expression ratios and statistics. Differential expression was tested by Student’s *t*-test and corrected for multiple testing using a false discovery rate (FDR) *q*-value of 0.05 (48). A list of significant differentially expressed genes (log2 ratio > 1.0; FDR *q*-value = 0.05) was subject to “Signature Search” using Genevestigator (https://genevestigator.com/gv/) (53). Additional analyses were performed on the entire list of expressed genes using non-parametric GO analysis using the Gorilla website (http://cbl-gorilla.cs.technion.ac.il/) (54) and the program Genevestigator with default settings.

### qRT-PCR confirmation

qRT-PCR was performed similarly to Wallström et al. (2010) (55). Four independent RNA samples were analyzed out in triplicates using the MAXIMA First strand synthesis kit (K1671; Thermo Scientific Waltham, MA USA) using manufacture’s guidelines. The resulting cDNA was quantified by qPCR using the MAXIMA SYBR green kit with 40 cycles in a Rotor-gene Q thermocycler (Qiagen Venlo, Netherlands), following the settings: 95 °C, for 2 min; and cycles of (95 °C, 20 sec; 60°C, 15 s; 72°C, 15 s; 81°C, 20 s); 1 cycle of 50°C, 1 min. Signals were acquired at 72°C and 81°C for analysing consistency in accumulation, and the comparative quantification was performed on data acquired at 81 °C.. Three technical replicates were run for each biological replicate. Gene products of the housekeeping gene *UPL7* (At3g53090) and three experimental genes *NIT1* (At3g44310), *PP2AA3* (At1g13320) and *EXP5* (At3g29030) were analysed (S7). Each reaction had a volume of 20 µL, and contained cDNA corresponding to 0.16 ng of DNase-treated RNA. Product identity and specificity was confirmed by melt curve analysis, as well as gel electrophoresis of representative and ambiguous samples. Individual reactions that showed unspecific products were deleted from analysis. Primers are listed in S8.

## Data availability

The RNA-Seq data in this study was deposited to the NCBI GEO under the accession number PRJNA573628. Figures, datasets and supplemental information have associated raw data that supported the findings of this study and that are available from the corresponding author upon reasonable request.

## Author contributions

A.G.R., P.G.B., and K.F. conceived the project. B.R.D. wrote the manuscript and contributed towards all molecular and genetic experiments with *Arabidopsis*, and contributed to analysis of *Streptomyces* volatiles. V.V. conducted the majority of *Streptomyces* volatile collection and analysis. All authors contributed towards editing of the manuscript.

## Additional information

For all correspondence please contact A.G.R. or P.G.B.

## Acknowledgements

This project was supported by Plant link, the Crafoord foundation, ICE-3 (SLU, Formas), the SLU Centre for Biological Control (CBC). We would like to thank Dr. G. Eric Schaller at Dartmouth College for donation of the *kmd1,2,4* and *kmd1,2,α4* seed stock and Dr. Matthew Escobar at California State University at San Marcos for the seed stocks of *arr1,10,12* and DR5::GFP. We are also thank Dr. Olivier van Aken at Lund University for critical reading of the manuscript and Dr. Oscar M. Rollano Peñaloza at Lund University for advice with transcriptomic analysis.

