## Supplemental Information for "The *Streptomyces* volatile 3-octanone alters auxin/cytokinin and growth in *Arabidopsis thaliana* via the gene family *KISS ME DEADLY*"

S1 | **Growth data for Col-0 response to gas-phase connection to different amounts of each *Streptomyces* strain.**

| Experimental conditions | LRG; LRG (cm) $day^{-1}$ $LR^{-1}$ | PRG; PRG (cm) $day^{-1}$ $PR^{-1}$ | log2 PRG/LRG | LRE; LRE (count) $day^{-1}$ $PR^{-1}$ |
|---|---|---|---|---|
| Control | 0.19 ± 0.004 | 0.97 ± 0.01 | 2.35 ± 0.03 | 3.91 ± 0.11 |
| 2 $cm^2$ *S.c.* M145 | 0.22 ± 0.02 | 0.77 ± 0.05 | 1.81 ± 0.03 | 3.66 ± 0.04 |
| 5 $cm^2$ *S.c.* M145 | 0.25 ± 0.01 | 0.64 ± 0.04 | 1.35 ± 0.07 | 3.79 ± 0.11 |
| 10 $cm^2$ *S.c.* M145 | 0.40 ± 0.01 | 0.34 ± 0.01 | -0.25 ± 0.03 | 3.74 ± 0.13 |
| 2 $cm^2$ *S.c. bldA* | 0.26 ± 0.03 | 0.81 ± 0.08 | 1.62 ± 0.10 | 3.79 ± 0.06 |
| 5 $cm^2$ *S.c. bldA* | 0.42 ± 0.03 | 0.77 ± 0.06 | 0.86 ± 0.03 | 4.19 ± 0.23 |
| 10 $cm^2$ *S.c. bldA* | 0.31 ± 0.02 | 0.18 ± 0.02 | -0.78 ± 0.08 | 3.89 ± 0.12 |
| 2 $cm^2$ *S.c. bldM* | 0.23 ± 0.03 | 1.00 ± 0.04 | 2.13 ± 0.22 | 3.11 ± 0.39 |
| 5 $cm^2$ *S.c. bldM* | 0.38 ± 0.03 | 0.44 ± 0.01 | 0.56 ± 0.11 | 3.56 ± 0.24 |
| 10 $cm^2$ *S.c. bldM* | 0.46 ± 0.02 | 0.51 ± 0.02 | 0.13 ± 0.07 | 4.23 ± 0.08 |
| 2 $cm^2$ *S.c. geoA mibAB* | 0.21 ± 0.02 | 1.06 ± 0.06 | 2.33 ± 0.22 | 4.31 ± 0.37 |
| 5 $cm^2$ *S.c. geoA mibAB* | 0.38 ± 0.02 | 0.73 ± 0.02 | 0.94 ± 0.06 | 4.28 ± 0.39 |
| 10 $cm^2$ *S.c. geoA mibAB* | 0.29 ± 0.01 | 0.22 ± 0.02 | -0.40 ± 0.09 | 3.76 ± 0.07 |
| 2 $cm^2$ *S.v.* | 0.21 ± 0.01 | 0.88 ± 0.02 | 2.05 ± 0.08 | 3.50 ± 0.06 |
| 5 $cm^2$ *S.v.* | 0.21 ± 0.01 | 0.81 ± 0.01 | 1.92 ± 0.08 | 3.43 ± 0.07 |
| 10 $cm^2$ *S.v.* | 0.18 ± 0.01 | 0.85 ± 0.01 | 2.22 ± 0.05 | 3.37 ± 0.03 |

Seven-day-old Col-0 seedlings where exposed for two days to volatiles emanating from differently sized plates containing *S. coelicolor* (*S.c.*) genotypes or *S. venezuelae* (*S.v.*). Growth rate was calculated as root length extension for control and volatile-exposed seedlings. Lateral Root Growth (LRG) per Lateral Root (LR), Primary Root Growth (PRG) per Primary Root (PR), the log2 ratio between PRG and LRG, (PRG/LRG) and Lateral Root Emergence (LRE) per PR are given ± standard error.

S2 | **Growth of *A. thaliana* Col-0 in response to *Streptomyces* volatiles.** Seven-day-old seedlings of *A. thaliana* Col-0 were treated ± gas-phase contact with 10 $cm^2$ *Streptomyces* agar cultures for six additional days. **a**, Control growth. **b**, *S. coelicolor* (*S.c.*) M145 wild-type on SFM medium. **c**, *S. venezuelae* (*S.v.*) grown on MYM medium. **d-e** Developmental *S. coelicolor* mutants *bldA* (**d**) and *bldM* (**e**) cultured on SFM. **f**, *S. coelicolor* mutant *geoA mibAB* cultured on SFM. **g-h**, seedlings grown on plant media without sucrose and treated ± volatiles from M145 cultured on SFM. **i-j**, Seedlings grown as in **a-f** and treated with volatiles from M145 cultured on MYM (**i**) and *S. venezuelae* cultured on SFM (**j**).

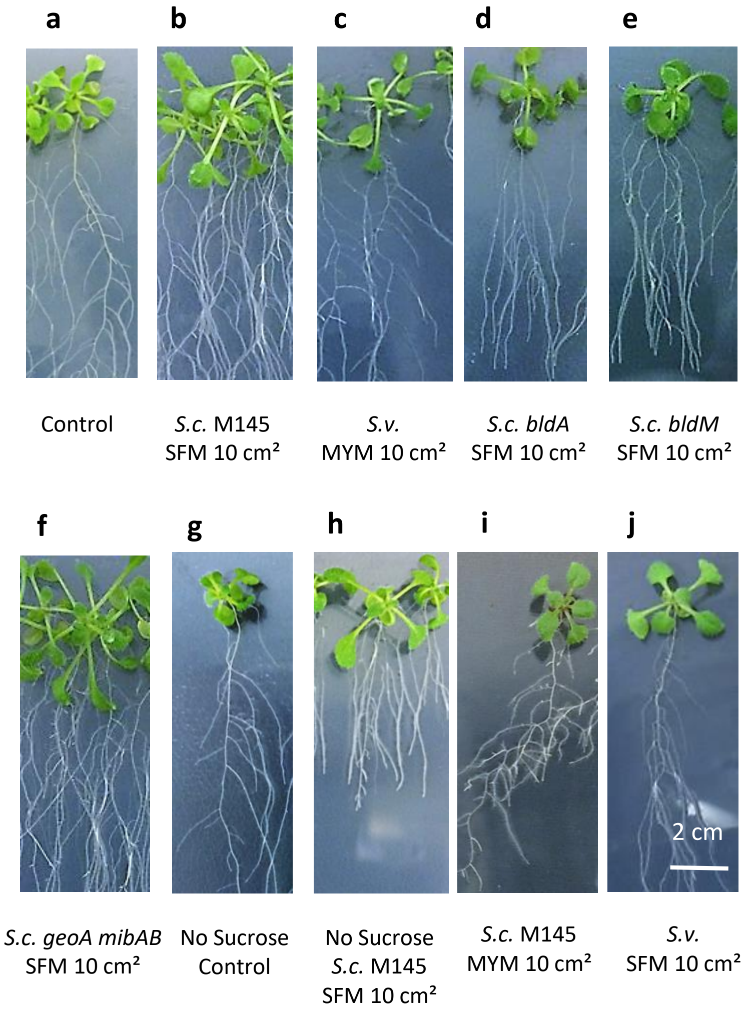

S3 | **Growth of *A. thaliana* hormone mutants in response to *S. coelicolor* M145 volatiles.** Seven-day-old seedlings of *A. thaliana* hormone mutants were treated ± volatiles from 10 cm² of *S. coelicolor* M145 culture on SFM for additionally six days.

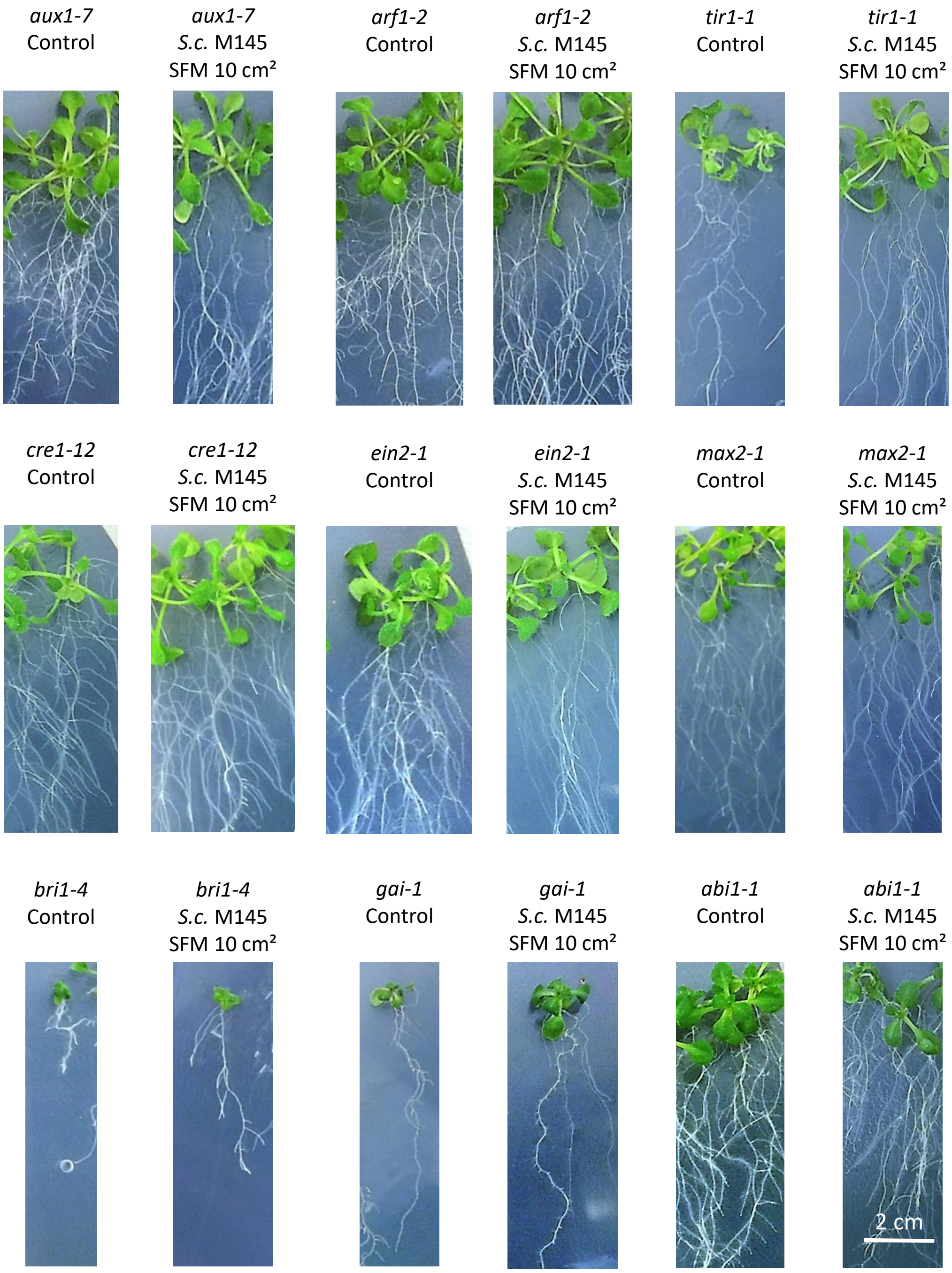

S4 | **DR5::GFP response to *S. coelicolor* volatiles.** Representative figures of primary root tips of the DR5::GFP line treated over different times ± *S. coelicolor* volatiles are shown. Seven-day-old DR5-GFP seedlings were exposed for 2, 6 and 24 h to volatiles from *S. coelicolor* M145 grown on 10 $cm^2$ SFM medium or to control plates. Roots were analyzed by bright field and fluorescence microscopy using a GFP-filter (excitation at 457 - 487 nm, emission 502 - 538 nm) coupled to a Nikon-Optiphot-2 microscope (Nikon Corporation, Tokyo, Japan). DR5::GFP expression visualize a redistribution of the auxin response upon exposure to *S. coelicolor* volatiles. DR5::GFP shows little difference in fluorescence distribution after 2 h of *S. coelicolor* volatiles as compared to control. However, after 6 h treatment, an increase in fluorescence could be observed in the vascular tissue relative to the apical meristem (*cf.* arrowheads) and after 24 h a general tissue-wide diffuse fluorescence was specifically observed in the volatile-treated root (brackets).

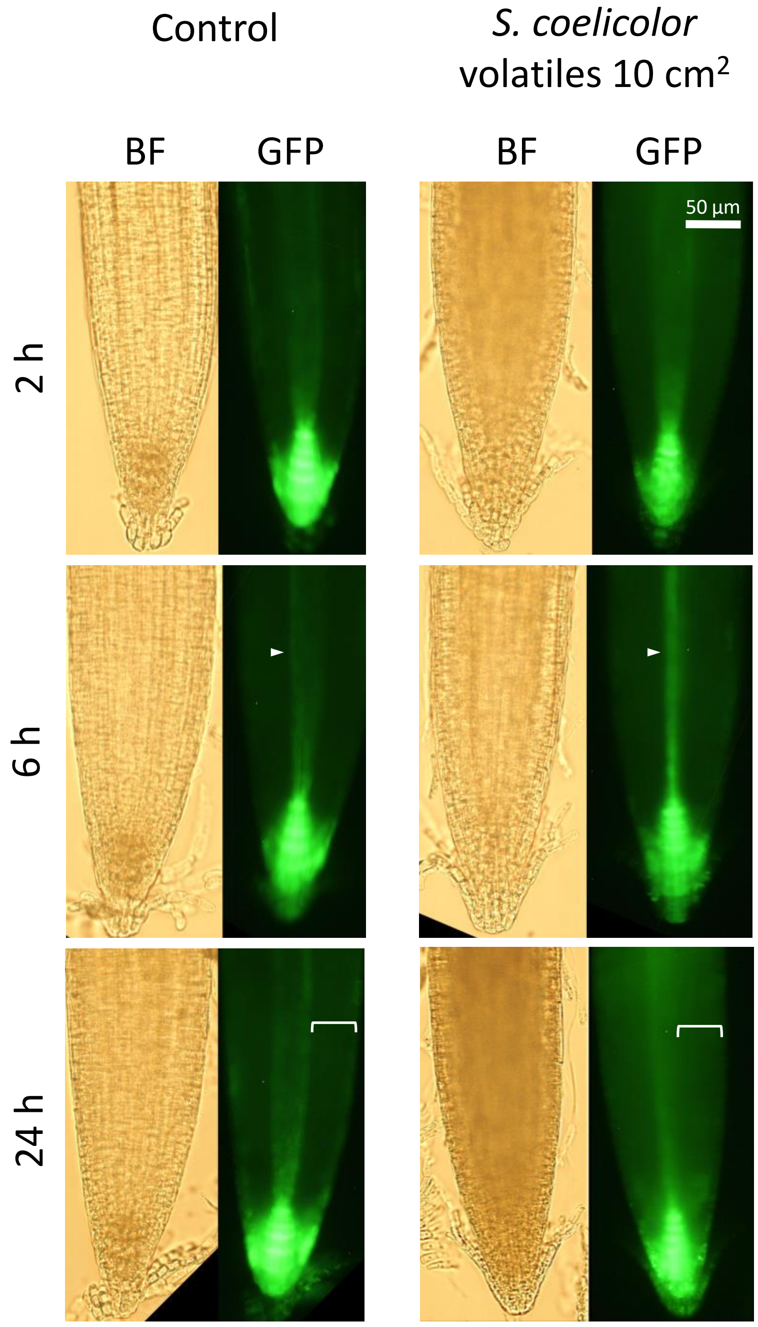

S5 | **Growth of *A. thaliana* mutants for *KMD*-associated genes in response to *S. coelicolor* volatiles.** The images show representative 7-day-old seedlings of *A. thaliana KMD* and associated cytokinin hormone mutants that have been treated ± 10 cm² of *S. coelicolor* M145 volatiles for an additional six days.

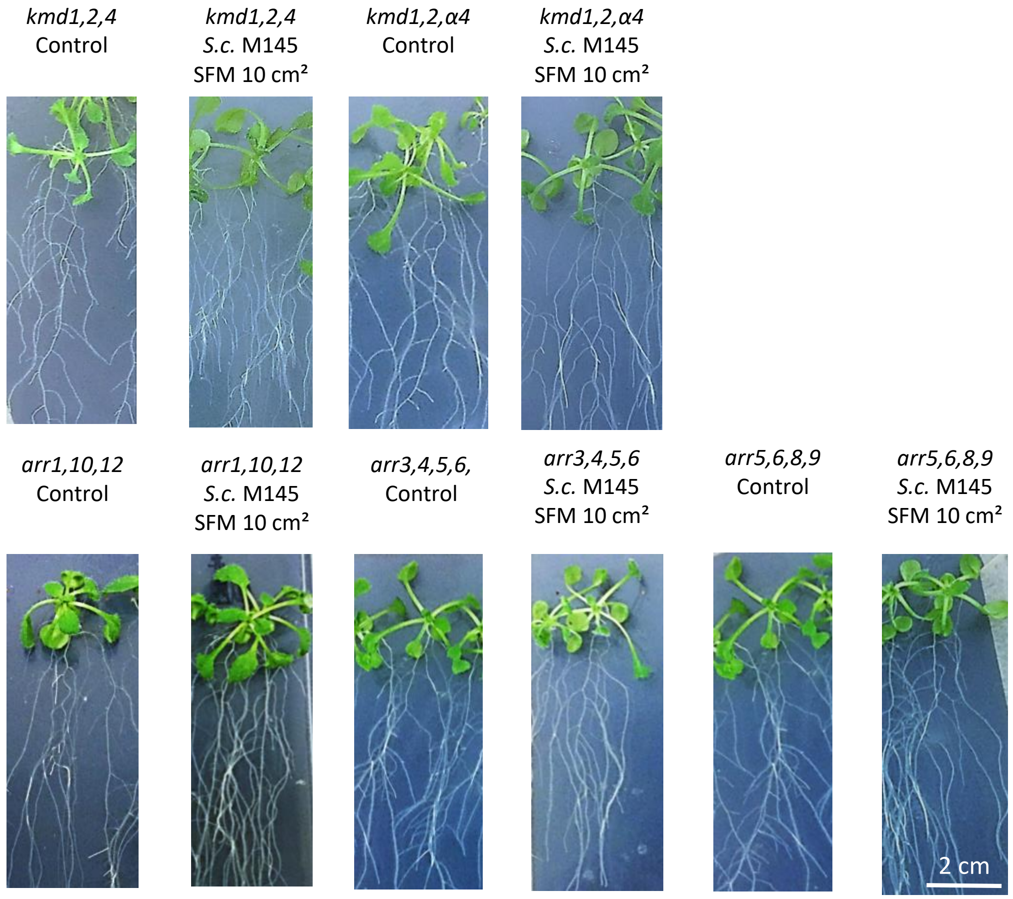

S6 | **Genetic stocks.** Genotypes of *Arabidopsis thaliana* seed lines and *Streptomyces* strains used in this publication.

| ***Name*** | ***Gene*** | ***Ecotype*** | ***References*** |
|---|---|---|---|
| Col-0 | n.a. (Wild-type) | Col-0 | n.a. |
| *arf1-2* | At1g59750 | Col-0 | Okushima *et. al.* 2005 |
| *arr1-3,10-1,12-1* | At3g16857, At4g31920, At2g25180 | Col-0 | Ishida *et. al.* 2008 |
| *arr3-1,4-1,5-1,6-1* | At1g59940, At1g10470, At3g48100, At5g62920 | Col-0 | To *et. al.* 2004 |
| *arr5-1,6-1,8-1,9-1* | At3g48100, At5g62920, At2g41310, At3g57040 | Col-0 | To *et. al.* 2004 |
| *aux1-7* | At2g38120 | Col-0 | Pickett *et. al.* 1990 |
| *cre1-12* | At2g01830 | Col-0 | Higuchi *et. al.* 2004 |
| DR5::GFP | pDR5::GFP (construct) | Col-0 | Ottenschlager *et. al.* 2003 |
| *ein2-1* | At5g03280 | Col-0 | Guzman and Ecker 1990 |
| *etr1-1* | At1g66340 | Col-0 | Bleecker *et. al.* 1988 |
| *kmd1-1,2-1,4-1* | At1g80440, At1g15670, At3g59940 | Col-0 | Kim H. *et al.* 2013 |
| *kmd1-1,2-1,α4* | At1g80440, At1g15670, (At2g44130), At3g59940 | Col-0 | Kim H. *et. al.* 2013 |
| *max2-1* | At2g42620 | Col-0 | Stirnberg *et. al.* 2002 |
| *tir1-1* | At3g62980 | Col-0 | Ruegger *et. al.* 1998 |
| *abi1-1* | At4g26080 | Ler-0 | Koornneef *et. al.* 1984 |
| *gai-1* | At1g14920 | Ler-0 | Koornneef *et. al.* 1985 |
| *bri1-4* | At4g39400 | Ws-2 | Noguchi *et. al.* 1999 |
| *coi1-37* | At2g39940 | Ws-2 | Kim J. *et. al.* 2013 |

| ***Strains*** | ***Relevant Gene Mutations*** | ***Species and Parental Strain*** | ***Source or Reference*** |
|---|---|---|---|
| M145-(*S.c.*) | Prototrophic, SCP1$^{-}$ SCP2$^{-}$ | *Streptomyces coelicolor*A3(2)-(M145) | Kieser *et. al.* 2000 |
| J2192-(*geoA mibAB*) | Δ*geoA* Δ*mibAB::apr* | *Streptomyces coelicolor*A3(2)-(M145) | Buttner & Bibb (JIC) |
| J1681-(M600) | Prototrophic, SCP1$^{-}$ SCP2$^{-}$ | *Streptomyces coelicolor*A3(2)-(M600) | Kieser *et. al.* 2000 |
| J1681-(*bldA*) | Δ*bldA::apr* | *Streptomyces coelicolor*A3(2)-(M600) | Kim *et. al.* 2005 |
| J3445-(*bldM*) | Δ*bldM::apr* | *Streptomyces coelicolor*A3(2)-(M600) | Kieser *et. al.* 2000 |
| NRRL B-65442-(*S.v.*) | Wild-type | *Streptomyces venezuelae*-(NRRL B-65442) | Bush *et. al.* 2019 |

S7 | **qRT-PCR verification of RNA-Seq expression.** Comparison of transcript ratios between 7-day-old Col-0 seedlings that have exposed to volatiles from a 10 $cm^2$ culture of *S. coelicolor* M145 for 2 h and  control seedlings.. Comparison of expression values was made by qRT-PCR and RNA-Seq. Each gene signal was normalized to the *UBIQUITIN PROTEIN LIGASE7* (At3g53090) internal control gene in the qRT-PCR analysis. Bars represent standard errors.

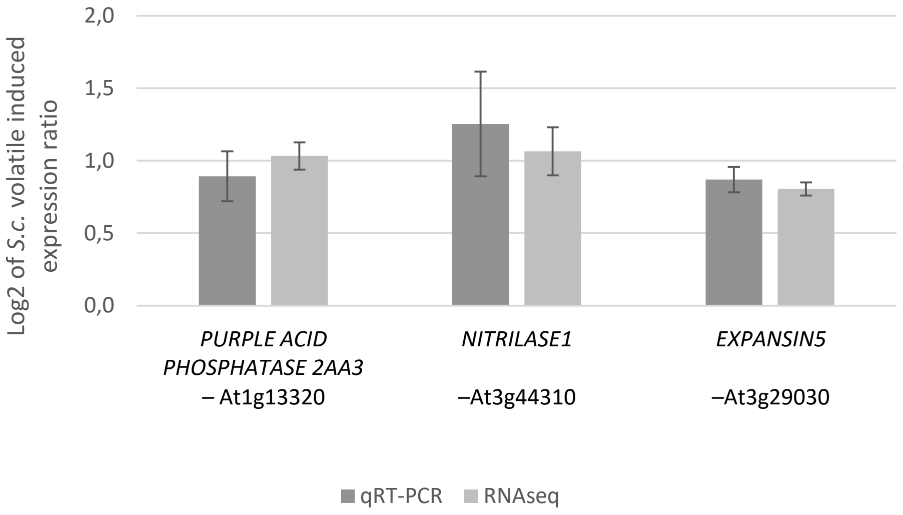

S8 | **Primer sequences**. Gene targets, forward and reverse primer sets and product sizes for qRT-PCR verification.

| Target Gene | Forward | Reverse | Product Size |
|---|---|---|---|
| *UPL7* | TGCTACAATACTCTTAAGCTTCCAACG | GTGCATAACAAGATGAATACCTGGTT | 136 |
| *PP2AA3* | ACCAGCTGAAAGTCGCTTAGC | GCTATGGCGGAAGAGTTGGG | 213 |
| *EXP5* | CCGCATGCTCCACCCATAG | CGCTTCTCGTGGTTCATCTCC | 160 |
| *NIT1* | TCGTCACAGCTGATATTGATATAGC | TCCTCGGGTGCTCATTTACGG | 142 |
